# A shared neural architecture underlies finger movement encoding in the human sensorimotor cortex

**DOI:** 10.1101/2025.09.11.675501

**Authors:** Theo Marins, Frederico Augusto Casarsa de Azevedo, Fernanda Tovar-Moll, Guilherme Wood

## Abstract

Human behavior evokes cortical responses that are highly variable across individuals, challenging the existence of a shared informational architecture, previously reported in animal studies. Here, using 7-Tesla fMRI and Procrustean transformation, we show that high-dimensional alignment of neural information from finger-tapping sequences reveals a latent neural architecture for sequential finger movements that generalizes across brains. This shared representation is instantiated in both sides of the sensorimotor cortex and does not reflect trial-wise variability of motor behavior, as regional decoding differences persisted after controlling for movement-time fluctuations. This conserved sensorimotor encoding structure provides a neuroscientific foundation for scalable, calibration-free brain-computer interfaces and cross-subject models of motor rehabilitation.

## Introduction

Finger-tapping movements are a popular paradigm for investigating sensorimotor representations in the human brain. To execute these movements, a combination of multiple parameters, such as velocity, joint angles, and spatial position, needs to be at play. Given that the brain’s topographic surface structure varies greatly across individuals, the brain responses elicited by sequential finger movements are perceived as highly idiosyncratic^1–3^.

Animal studies have shown that a shared neural representation space for voluntary movements exists across individuals within the same species^4,5^, suggesting that the evolutionary constraints that have preserved the overall neural circuits across individuals^6^ have also impacted how the neural information is instantiated in the brain. In humans, it remains to be shown whether individual brains represent sequential finger movements in idiosyncratic ways or whether common evolutionary forces constrain the overall neural dynamics to preserve a neural representation.

Previous human studies showed the feasibility of the high-dimensional alignment of sensorimotor activity to investigate these concepts^7,8^, suggesting a compelling direction for future research. However, it is still unclear to what extent these previous findings truly captured shared information encoded in idiosyncratic cortical topographies or reflected uncontrolled sources of structured variance, including violations of noise assumptions inflating false positives^9,10^, insufficient control of multiple comparisons^11,12^, or systematic behavioral variability shaping trial-wise BOLD responses^13,14^. Using high-field (7-Tesla) brain mapping of finger-tapping sequences, we investigated the existence of a reliable shared neural code of movements in the human brain. By avoiding the confounding effects of earlier studies, we demonstrate cross-subject generalization in a brain-decoding model. Our approach reveals a shared high-dimensional representational architecture for encoding finger movements; we also describe its spatial extent in the sensorimotor cortex.

The finding of cross-subject latent neural dynamics underlying complex behavior has the potential to impact both the theoretical frameworks of neural computation and applied neuroscience, as it enables better alignment strategies that yield brain decoders that can be trained on one participant and translated to other individuals^15,16^. This possibility allows a brain-computer interface to be pre-programmed from healthy volunteers and smoothly applied to clinical populations without lengthy customized calibration. Ultimately, a shared neural code is the foundation for scalable, widely accessible neurotechnology afield.

## Results and Discussion

For eight runs, 12 right-handed subjects performed 12 unique finger-tapping sequences, which were matched by their difficulty level and starting finger, number of finger repetitions in a sequence, and first-order finger transitions^17^. Each unique finger-tapping sequence consisted of two repetitions of nine-digit movements that were performed at the same pace with the five fingers of the right hand (Figure 1A, numbered squares), and repeated thrice per run. To avoid bias, we analyzed the first pair of perfectly executed movements of each sequence per run, excluding the remaining pairs of trials (Figure 1A, gray squares). Individual brain response patterns for the selected pair of trials were obtained with a general linear model (GLM; see Methods), and the resulting z-scores were used for subsequent analyses. For each side of the sensorimotor cortex, defined using the Destrieux Atlas^18^, all runs from one participant were held out as test data, while data from all remaining participants were used as the training data (Figure 1B). Next, we applied Procrustean transformations to the training data within each surface-based searchlight to align individual brain response patterns in a common, high-dimensional space^19,20^, and to train the decoding model (linear support vector machine (SVM) with L2 regularization). The test data were then projected into the common space and used to test the decoding model. This procedure was repeated so that each participant served as the test subject once.

**Figure 1.**
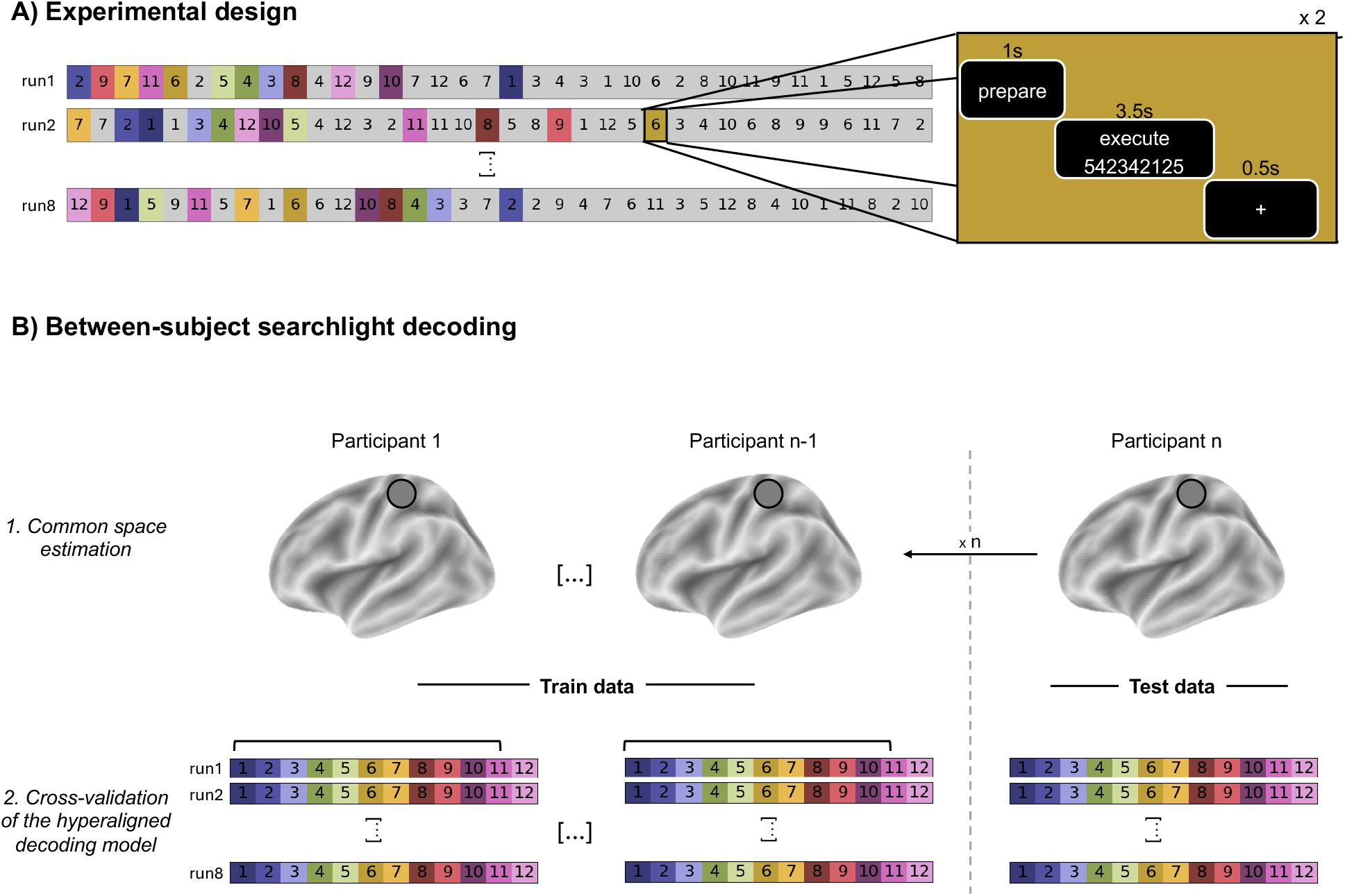
Experimental design and the between-subject searchlight decoding of hyperaligned brain data. (A) Each of the twelve sequences of finger-tapping movements was performed twice in a row, over three times in each of the eight runs. The first pair of perfectly executed finger-tapping sequences (colored squares) was modeled using a general linear model (see Methods), and the resulting z-scored map was used for subsequent analysis. (B) In a leave-one-subject-out scheme, the common space was estimated within each surface searchlight based on the neural representations of all but one subject (training data), then the brain data from the left-out subject (test data) was transformed into the common space (1. Common space estimation). Finally, the training and test data were used to train and test the linear classifier, respectively (2. Cross-validation of the hyperaligned decoding model). This process was repeated 12 times (n) so that each subject served as a test subject once, testing the classifier’s generalization across subjects.

The statistical significance of the decoder performance was assessed using non-parametric permutation testing. For each searchlight vertex and test subject, the empirical null distribution was estimated by repeating the classification 100 times with randomly permuted sequence identity labels in the training data. Group-level inference was performed by aggregating subject-level accuracies across test subjects, and empirical p-values were computed as the proportion of null distribution values exceeding the observed group mean accuracy (with a +1 correction), corresponding to a one-sided test for above-chance decoding. Resulting p-values were corrected for multiple comparisons using the Benjamini–Hochberg false discovery rate (FDR) procedure^21^.

Here, we show that behind the apparent highly idiosyncratic brain response elicited by sequential finger movements lies a latent informational architecture shared across participants. Based on permutation-based statistics, we observed above-chance-level accuracy in sensorimotor cortical areas in both hemispheres (Figure 1, Table 1), corroborating the notion that the sensorimotor cortex in both hemispheres contains information about unimanual sequential finger movements^22^.

**Table 1.**

| Label | Left | Right |
| --- | --- | --- |
| Central Sulcus | 60.97 (1.18) | 60.06 (1.04) |
| Postcentral Gyrus | 47.29 (0.79) | 46.92 (0.85) |
| Postcentral Sulcus* | 54.4 (1.14) | 57.2 (1.06) |
| Precentral Gyrus* | 48.12 (0.88) | 46.79 (0.75) |
| Precentral Sulcus (inf.)* | 49.64 (1.0) | 41.96 (0.86) |
| Precentral Sulcus (sup.) | 53.75 (1.0) | 54.08 (1.02) |

Mean (and standard error) decoding accuracy by cortical label and hemisphere.
\*Cortical labels that showed the main effect of side (corrected for false discovery rate, $p < 0.05$ ). inf: inferior; sup: superior.

In brain decoding, higher decoder scores suggest highly stable and robust neural codes^23,24^. Thus, we compared accuracy scores using a repeated-measures ANOVA (within-subjects factors: cortical label and hemisphere) to investigate sensorimotor areas in which the representational geometry of neural patterns is more consistent across individuals. The central sulcus displayed the greatest decoding scores (main effect of cortical label, Greenhouse– Geisser–corrected: *F*(5, 55) = 347.7, *p* < .001, η^2^_p_ = 0.97, pairwise t-test with FDR-corrected *p* < 0.05), in line with previous evidence showing that this region robustly encodes sequential finger movement features, such as finger movements^22^, sequence identity^17^, and their preparation and execution^25^ in within-subject designs. A left-lateralization, contralateral to the moving fingers, was also detected (main effect of hemisphere: *F*(1, 11) = 18.35, *p* = .001, η^2^_p_ = 0.63; t-test: t(11) = 4.28, p = 0.001, Cohen’s d = 0.38). Finally, the central sulcus showed the highest decoder accuracy within each hemisphere (interaction between cortical label and hemisphere, Greenhouse–Geisser–corrected: *F*(5, 55) = 49.96, *p* < .001, η^2^_p_ = 0.82, pairwise t-test with FDR-corrected *p* < 0.05; Table 2).

**Table 2.** Left hemisphere

| Labels |  | T | p-corr |
| --- | --- | --- | --- |
| Central Sulcus | Postcentral Gyrus | 23.50 | < 0.001 |
| Central Sulcus | Postcentral Sulcus | 11.84 | < 0.001 |
| Central Sulcus | Precentral Gyrus | 25.94 | < 0.001 |
| Central Sulcus | Precentral Sulcus (inf.) | 15.20 | < 0.001 |
| Central Sulcus | Precentral Sulcus (sup.) | 11.39 | < 0.001 |
| Postcentral Gyrus | Postcentral Sulcus | -14.25 | < 0.001 |
| Postcentral Gyrus | Precentral Sulcus (inf.) | -4.89 | < 0.001 |
| Postcentral Gyrus | Precentral Sulcus (sup.) | -12.46 | < 0.001 |
| Postcentral Sulcus | Precentral Gyrus | 12.36 | < 0.001 |
| Postcentral Sulcus | Precentral Sulcus (inf.) | 7.90 | < 0.001 |
| Precentral Gyrus | Precentral Sulcus (inf.) | -2.38 | 0.042 |
| Precentral Gyrus | Precentral Sulcus (sup.) | -14.19 | < 0.001 |
| Precentral Sulcus (inf.) | Precentral Sulcus (sup.) | -5.22 | < 0.001 |

**Right hemisphere**
| Labels |  | T | p-corr |
| --- | --- | --- | --- |
| Central Sulcus | Postcentral Gyrus | 48.7172 | < 0.001 |
| Central Sulcus | Postcentral Sulcus | 8.0746 | < 0.001 |
| Central Sulcus | Precentral Gyrus | 25.4148 | < 0.001 |
| Central Sulcus | Precentral Sulcus (inf.) | 28.4256 | < 0.001 |
| Central Sulcus | Precentral Sulcus (sup.) | 8.7164 | < 0.001 |
| Postcentral Gyrus | Postcentral Sulcus | -25.3801 | < 0.001 |
| Postcentral Gyrus | Precentral Sulcus (inf.) | 9.2577 | < 0.001 |
| Postcentral Gyrus | Precentral Sulcus (sup.) | -11.0499 | < 0.001 |
| Postcentral Sulcus | Precentral Gyrus | 21.5220 | < 0.001 |
| Postcentral Sulcus | Precentral Sulcus (inf.) | 24.7336 | < 0.001 |
| Postcentral Sulcus | Precentral Sulcus (sup.) | 4.8094 | < 0.001 |
| Precentral Gyrus | Precentral Sulcus (inf.) | 9.1187 | < 0.001 |
| Precentral Gyrus | Precentral Sulcus (sup.) | -15.1928 | < 0.001 |
| Precentral Sulcus (inf.) | Precentral Sulcus (sup.) | -14.0480 | < 0.001 |

Pairwise posthoc t-tests of decoder accuracy per cortical label within the left (top) and right (bottom) hemispheres. Only statistically significant results (corrected for false discovery rate, $p < 0.05$ ) are depicted. inf: inferior; sup: superior.

Although participants were instructed to perform the sequential finger movements at a fixed pace^17^, residual variability was observed (coefficient of variation: 16.34 ± 6.59%). Therefore, we tested whether trial-to-trial differences in motor execution could account for variability in decoding accuracy, as behavioral fluctuations can introduce structured variance in the BOLD response that may spuriously inflate classification performance^13,14^. To this end, we fitted a linear mixed-effects model to assess whether regional decoding differences persisted after controlling for movement-time variability. Even though there was a significant probability that movement-time variability affected accuracy (p = 0.001, 95% CI = [0.289, 1.059]), its effect was smaller (β = 0.67 per standard deviation) than that of large regional differences (β ≈ −13 to −15). Importantly, regional effects remained robust after controlling for movement-time variability (all |β| > 6, p < 0.001), indicating that movement-time variability alone cannot explain the observed cortical pattern.

We also analyzed sequence encoding using anatomical alignment as a reference, following the same procedure as for the hyperalignment data, except that participants’ functional data remained on the template surface fsaverage. Under anatomical alignment, decoding accuracy mean values were close to chance; 8.59% (0.02% ± SE) and 8.32% (0.03% ± SE) for the left and right hemispheres, respectively. In contrast, the hyperaligned decoding model improved the mean accuracy across 100% of the sensorimotor cortex vertices; 53.06% (0.21% ± SE) and 52.29% (0.24% ± SE) in the left and right hemispheres, respectively. These findings strongly indicate the existence of a high-dimensional latent information space that is shared across individual brains^19,26^.

Besides the technical feasibility of applying hyperalignment to the motor system^7,8^, we show that the information it captures reflects the genuine neural structure rather than methodological artifacts. Our findings suggest that neural codes for complex sequential finger movements are topographically conserved across individuals, enabling cross-subject decoding within the human sensorimotor cortex.

Currently, applications of clinically-oriented brain decoding requiring user-specific calibration are extremely time consuming^27^, mainly due to the high inter-subject/session variability^28^. Our findings provide compelling evidence that despite individual variability in cortical topography, the human brain expresses a shared, high-dimensional encoding architecture for sequential finger movements. This shared neural code lies the groundwork for future work in personalized neurotechnologies, scalable brain-computer interfaces, and cross-subject models of motor learning and rehabilitation.

## Acknowledgments

This work was supported by the Austrian Science Fund FWF (doi: 10.55776/ESP397), the Foundation for Research Support in the State of Rio de Janeiro (FAPERJ), and intramural grants of the D’Or Institute for Research and Education (IDOR). We thank Marcia Triunfol for the insightful comments on the manuscript.

## Materials and Methods

Here, we analyze a dataset available in OpenNeuro repository, published elsewhere^17^. The experimental description below was obtained from publicly available sources, such as the original publication^17^ and the dataset description. Specific parts irrelevant to the present study (e.g., motor learning procedure and other fMRI sessions) have been omitted for clarity purposes. Refer to the original publication for a detailed methodological description.

### Participants

The high-field fMRI data used in this study are a publicly available dataset previously published by Berlot and colleagues^17^. We used fMRI data from 12 right-handed participants (8 females, mean age of 22.4, sd=2.84, sample details in SI). Informed consent and data usage agreement have been obtained from participants prior to the study. In it, it was emphasized that participants could withdraw from the study at any timepoint. The experimental procedures have been approved by the local ethics committee at Western University^17^, where the data have been collected.

### Data collection

MRI data were acquired using a 7-Tesla Siemens Magnetom scanner with a 32-channel receive head coil (8-channel parallel transmit). Anatomical T1-weighted scan used a magnetization-prepared rapid gradient echo sequence (MPRAGE) with a voxel size of 0.75 x 0.75 x 0.75 mm isotropic (field of view = 208 x 157 x 110 mm [A-P; R-L; F-H], encoding direction coronal). For fMRI data, the GRAPPA3 sequence (multiband acceleration factor 2, repetition time [TR]=1.0 s, echo time [TE]=20 ms, flip angle [FA]=30 deg, 44 slices with isotropic voxel size of 2 x 2 x 2 mm) has been used. A gradient echo field map has been collected to estimate magnetic field inhomogeneities (transversal orientation, field of view 210 210 160 mm and 64 slices with 2.5 mm thickness, TR = 475 ms, TE = 4.08 ms, FA = 35 deg).

### Experimental design

The dataset involved 26 participants performing 12 different finger-tapping movement sequences with their right hand, cued by visual stimuli, over eight runs^17^. Each sequence was repeated twice in a row, for a total of six times in every run. Participants were right-handed and had no prior history of psychiatric or neurological disorders.

The execution of each sequential finger movement began with a 1-s preparation time, during which the sequence was presented on a screen. After that time, a ‘go’ signal was displayed as a short pink and expanding line underneath the sequence numbers that indicated the speed at which participants were required to press along. The execution phase, including the feedback on the overall performance, lasted 3.5 s, and the inter-trial interval was 0.5 s. Each trial lasted 5 s in total. In each run, five resting periods of 10 s were added randomly between pairs of trials. For details about the experimental design, please refer to the original publication^17^.

We removed trials with execution errors, keeping data of the first pair of perfectly executed finger-tapping movements per participant, run, and sequence. This led to the exclusion of 14 participants, resulting in a sample of 12 subjects (8 females, mean age of 22.4, sd=2.84) who performed 12 sequential finger-tapping movements twice in a row, repeated three times per run (Figure 1).

### Data preprocessing

Functional and structural MRI data were preprocessed using fMRIPrep (version 23.2.1), a standardized and reproducible preprocessing pipeline. T1-weighted anatomical images were corrected for intensity non-uniformity, skull-stripped, and segmented into gray matter, white matter, and cerebrospinal fluid. Nonlinear spatial normalization to the MNI152NLin2009cAsym template was performed using brain-extracted T1-weighted images.

For each functional run, a reference BOLD image was generated and co-registered to the individual T1-weighted anatomical image using boundary-based registration with six degrees of freedom. Head motion parameters (three translations and three rotations) were estimated for each run, and framewise displacement (FD) was computed. No smoothing was performed. Preprocessed BOLD time series were projected onto the fsaverage5 cortical surface for each hemisphere for further analyses.

### Task events and trial selection

Task timing and condition information were extracted from run-specific event files. Each trial consisted of a finger-tapping sequence and was modeled with a fixed duration of 3.5 s. Sequence identity was defined by sequence number (1–12), with subject-specific remapping applied to ensure consistent labeling across experimental groups^17^. To minimize contamination from execution errors, trials were grouped into consecutive pairs of repetitions for each sequence. In each run, the first correctly executed pair of each finger-tapping sequence was selected, and the remaining were excluded from further analysis. At the subject-level, this resulted in eight pairs of executions per finger-tapping sequence. Each retained pair of sequences was treated as a unique experimental condition in the GLM.

### First-level surface-based GLM

First-level statistical analyses were performed using Nilearn’s First Level Model, implemented in surface space. Separate GLMs were fitted for each subject, run, and hemisphere. The design matrix included one task-related regressor per pair of sequences, convolved with the canonical SPM hemodynamic response function. The repetition time (TR) was set to 1 s. Nuisance regressors were included to account for non-neural sources of variance and comprised white matter signal, global signal, framewise displacement, and six rigid-body motion parameters. No temporal filtering, CompCor components, or autoregressive noise modeling were applied at the GLM stage beyond what is inherent to the design matrix specification. For each run-specific GLM, condition-specific contrasts were computed by isolating individual task regressors in the design matrix. This resulted in one surface-based z-statistic map per sequence-pair condition, per run, and per subject. Contrast maps were saved separately for cortical data and surface mesh geometry. The resulting z-maps were subsequently masked using a predefined sensorimotor cortex network^18^ after excluding nodes in the medial wall and served as inputs for the between-subject multivariate decoding analyses described below.

### Searchlight-based hyperalignment

To enable between-subject decoding of motor sequence identity, we implemented a surface-based searchlight hyperalignment procedure. For each hemisphere, overlapping cortical disks with a radius of 20 mm were defined on the fsaverage surface. Within each disk, neural response patterns from the training subjects were aligned to a common representational space using a Procrustes-based hyperalignment algorithm^29^.

A leave-one-subject-out cross-validation scheme was employed. In each fold, all runs from one participant were held out as test data, while data from all remaining participants were used to estimate the common model. The hyperalignment transformation was computed exclusively on the training data, ensuring that no test-subject information contributed to the construction of the common space. After estimating the common model within each searchlight disk, the test data were projected into the corresponding common representational space, enabling direct comparison of neural activity patterns across participants.

### Searchlight decoder implementation

Decoding of motor sequence identity was performed using a searchlight-based multivariate pattern classification approach^26^. Within each cortical disk of radius 10 mm, a linear SVM with L2 regularization was trained to classify neural activity patterns corresponding to the 12 finger-tapping sequences. Classification was performed on hyperaligned data, using training data from all non-test subjects and evaluating performance on the test data. Classification accuracy was computed for each searchlight and assigned to the central vertex, yielding one accuracy map per test subject and hemisphere. This procedure was repeated so that each participant served as the test subject exactly once.

### Anatomically-aligned decoder analysis

The between-subject decoding models with anatomically aligned data followed the same procedure as their counterparts on hyperaligned data, except that the individual functional data were kept in the template surface fsaverage.

### Permutation testing and group-level statistics of brain decoding maps

Statistical significance of decoding performance was assessed using non-parametric permutation testing. For each test subject and searchlight vertex, the decoding analysis was repeated 100 times with randomly shuffled sequence identity labels in the training data, while keeping the test labels unchanged. This yielded a subject-level null distribution of decoding accuracies for each vertex.

Group-level inference of the brain decoding maps was performed by aggregating observed decoding accuracies across all folds for each vertex. To estimate the empirical null distribution of group-level mean accuracy, the permutation-based accuracies were pooled across subjects. The empirical p-values were computed as the proportion of null distribution values greater than or equal to the observed group-level mean accuracy, with one added to both the numerator and denominator to reduce bias and avoid zero p-values^30^. This corresponds to a one-sided test for above-chance decoding. Resulting p-values were corrected for multiple comparisons across vertices using the Benjamini–Hochberg false discovery rate (FDR) procedure^21^. To improve visualization of the statistically significant (corrected FDR p-value < 0.05) findings, the plot displayed in Figure 2A was manually set.

**Figure 2.**
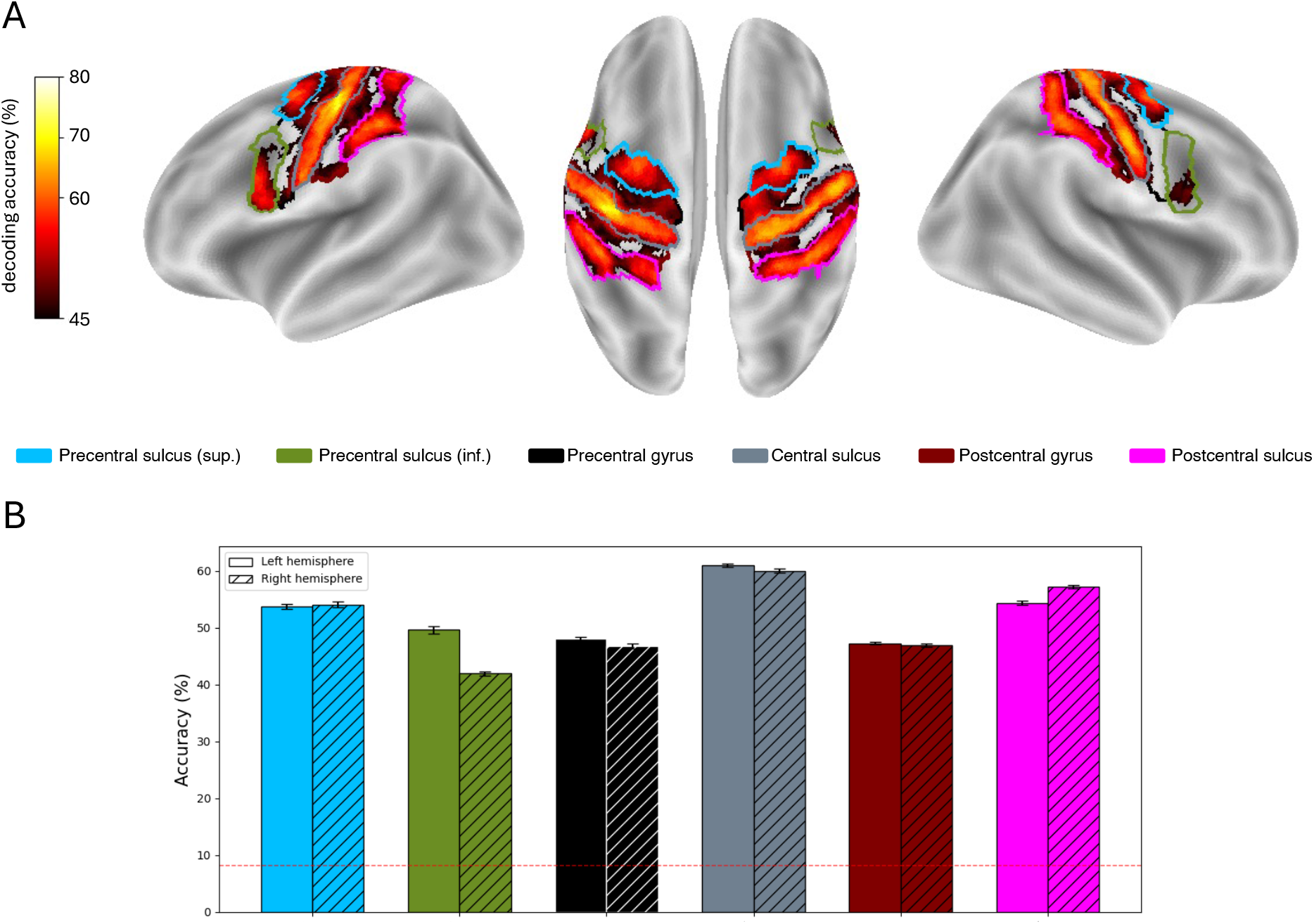
Group-level map of between-subject classification of sequence identity (sequences 1 to 12). (A) The heatmap shows vertex-wise classification accuracy obtained with permutation testing and false discovery rate (FDR) correction. (B) Average accuracy above chance level in each cortical label in the left and right hemispheres. Error bars indicate standard error. The theoretical chance level is 8.3% (red dashed lines). All cortical labels showed above chance-level accuracy. The sensorimotor cortical areas were obtained from the Destrieux Atlas^18^. Group statistics were performed based on subject-level data using a repeated-measures ANOVA (within-subjects factors: cortical label and hemisphere). We observed a main effect of cortical label (Greenhouse–Geisser–corrected: *F*(5, 55) = 347.7, *p* < .001, η^2^_p_ = 0.97), with the central sulcus showing the highest decoder accuracy (FDR-corrected *p* < 0.05). Moreover, a significant main effect of hemisphere (F(1, 11) = 18.35, p = .001, η^2^_p_ = 0.63), indicated a left-lateralization (t(11) = 4.28, p = 0.001, Cohen’s d = 0.38), contralateral to the moving fingers. Finally, a significant interaction between label and hemisphere (Greenhouse–Geisser–corrected: *F*(5, 55) = 49.96, *p* < .001, η^2^_p_ = 0.82) indicated that the central sulcus showed the highest decoder accuracy within each hemisphere (FDR-corrected *p* < 0.05).

**Figure 3.**
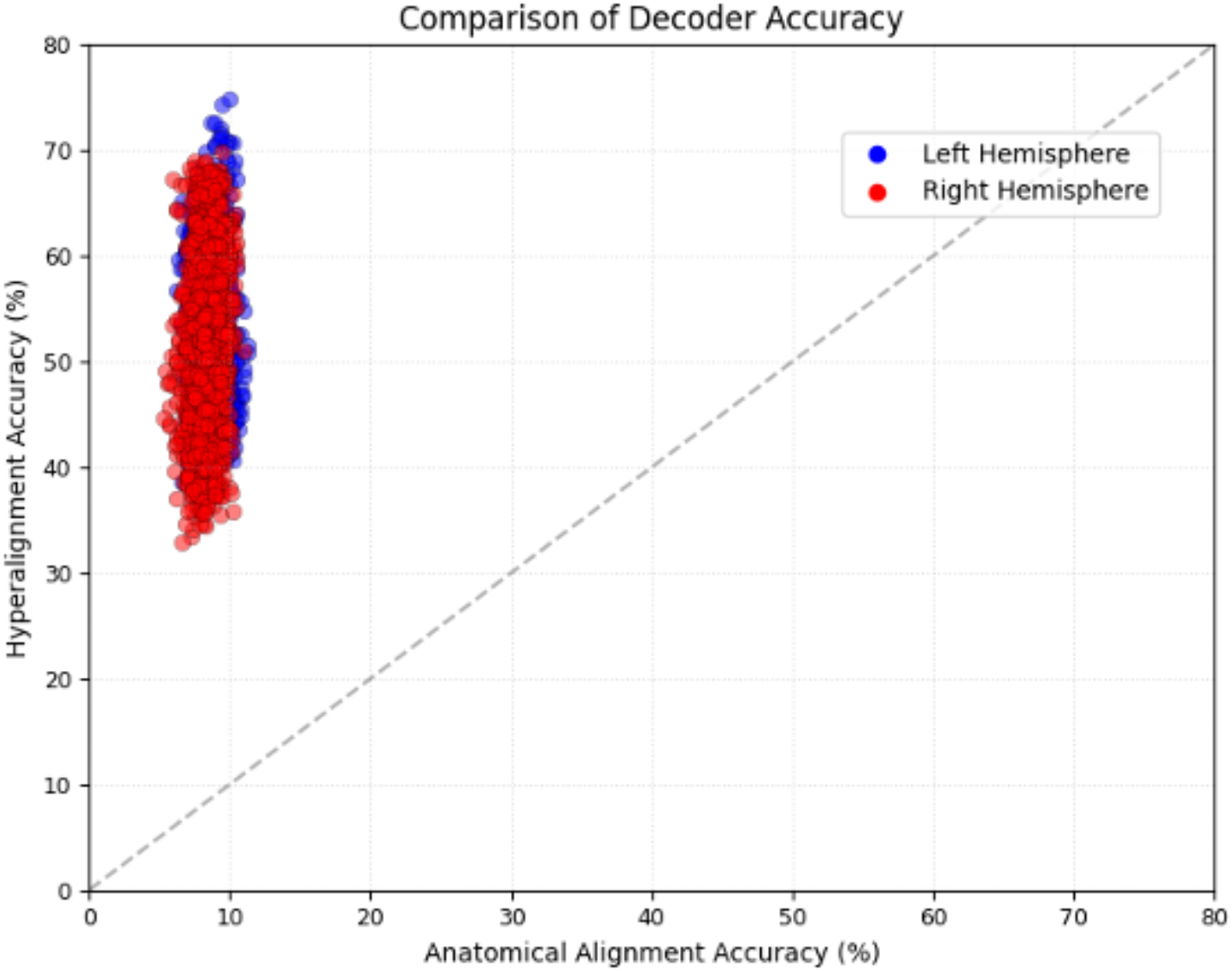
Comparison of decoder accuracy obtained with hyperaligned or anatomically aligned data, before false discovery rate correction for multiple comparisons. All vertices showed improved decoder accuracy with the hyperalignment model, and the mean absolute gains reached 44.5% and 44% on the left (blue dots, n=1275 vertices) and right (red dots, n=1204 vertices) hemispheres, respectively.

### Subject-level-derived statistics

Using the subject-level accuracy maps, we investigated the effects on cortical label and hemisphere. Cortical labels (six levels: precentral sulcus (sup), precentral sulcus (inf), precentral gyrus, central sulcus, postcentral gyrus and postcentral sulcus) and hemisphere (two levels: left and right) were compared in a repeated-measures ANOVA. Posthoc tests were performed to further investigate significant main effects, and FDR was implemented as the method for multiple comparison correction.

### Control analysis for movement-time variability

To test whether the variability in movement time accounted for differences in decoding accuracy, we fitted a linear mixed-effects model using restricted maximum likelihood estimation (REML). Accuracy was modeled as the dependent variable, with movement-time variability (z-scored), cortical region (Label), and hemisphere (Side) as fixed effects, including their interactions with movement time. Subject identity was included as a random intercept to account for repeated measures.

The model specification was: accuracy ∼ movement_time_std × (Label + Side) + (1 | subject_ID)

This approach allowed us to quantify the unique contribution of movement-time variability to accuracy while controlling for regional and hemispheric differences. This analysis directly tests whether regional accuracy differences persist after accounting for movement-time variability, thereby addressing the possibility that performance differences are driven by behavioral timing rather than neural representational structure.

### Data and code availability

fMRI data and analysis pipelines have been deposited to OpenNeuro (id ds002776). The codes used in the present study are available on GitHub (through GitFront): https://gitfront.io/r/theomarins/RFvFDXTnP875/cortical-motor-engram-decoding/.

